# Lack of astrocytic glycogen alters synaptic plasticity but not seizure susceptibility

**DOI:** 10.1101/2020.05.06.080978

**Authors:** Jordi Duran, M. Kathryn Brewer, Arnau Hervera, Agnès Gruart, Jose Antonio del Rio, José M. Delgado-García, Joan J. Guinovart

## Abstract

Brain glycogen is mainly stored in astrocytes. However, recent studies both *in vitro* and *in vivo* indicate that glycogen also plays important roles in neurons. By conditional deletion of glycogen synthase (GYS1), we previously developed a mouse model entirely devoid of glycogen in the central nervous system (GYS1^Nestin-KO^). These mice displayed altered electrophysiological properties in the hippocampus and increased susceptibility to kainate-induced seizures. To understand which of these functions is related to astrocytic glycogen, in the present study we generated a mouse model in which glycogen synthesis is eliminated specifically in astrocytes (GYS1^Gfap-KO^). Electrophysiological recordings of awake behaving mice revealed alterations in input/output curves and impaired long-term potentiation, similar, but to a lesser extent, to those obtained with GYS1^Nestin-KO^ mice. Surprisingly, GYS1^Gfap-KO^ mice displayed no change in susceptibility to kainate-induced seizures as determined by fEPSP recordings and video monitoring. These results confirm the importance of astrocytic glycogen in synaptic plasticity.

## 1 INTRODUCTION

Most cell types in the body store glucose in the form of glycogen, a branched macromolecule containing up to 55,000 glucose units. The only enzyme able to form glycogen *in vivo* is glycogen synthase (GYS). There are two isoforms of glycogen synthase in mammals: the muscle isoform, GYS1, which is expressed in all tissues except the liver, and the liver-specific isoform, GYS2. In the brain, glycogen is estimated to comprise about 0.1% of tissue weight [1]. Both astrocytes and neurons express GYS1 and synthesize glycogen, although glycogen levels in astrocytes are much higher than in neurons [2, 3].

In the past decades, numerous studies have demonstrated that brain glycogen plays a role in memory consolidation and synaptic function [reviewed in [4–6]]. In histological studies of the healthy brain, glycogen granules are almost always confined to astrocytic cell bodies and processes [7]. Hence, there is a longstanding belief that the contribution of brain glycogen to cerebral functions is entirely due to its role in astrocytes. However, recent *in vitro* studies suggested an active glycogen metabolism in neurons [2, 8].

To study the role of brain glycogen *in vivo*, we previously developed a transgenic mouse line (GYS1^Nestin-KO^) which lacked GYS1 and thus glycogen throughout the whole CNS, while GYS1 expression was normal in other tissues. Paired-pulse recordings at the CA3-CA1 synapse of the hippocampus showed that the GYS1^Nestin-KO^ animals displayed increased facilitation, i.e. an increased response to the second pulse [9, 10]. The GYS1^Nestin-KO^ animals also exhibited impaired long-term potentiation (LTP) evoked at the CA3-CA1 synapse. LTP is believed to be a primary molecular mechanism underlying long-term memory consolidation [9]. Additionally, the animals were more susceptible to hippocampal seizures induced by kainate or train stimulation [10]. These *in vivo* results demonstrate that brain glycogen plays a role in both short- and long-term synaptic plasticity as well as in the prevention of seizures.

Since the GYS1^Nestin-KO^ mice lacked both glial and neuronal glycogen, the differential contribution of each glycogen pool to these results could not be determined with this model. For this reason, we next generated a new model with greater cellular resolution devoid of GYS1 in a subset of glutamatergic neurons, the excitatory Ca^2+^/calmodulin-dependent protein kinase 2 (Camk2a)-positive neurons of the forebrain [11], including the pyramidal cells of the hippocampal CA3-CA1 synapse. Like the GYS1^Nestin-KO^, these animals presented altered LTP and an associative learning deficiency, although the impairment was not as pronounced. However, unlike the GYS1^Nestin-KO^ mice, GYS1^Camk2a-KO^ mice exhibited no statistically significant change in PPF and no difference in susceptibility to kainate-induced seizures. This study corroborated the presence of an active glycogen metabolism in neurons *in vivo* and illustrated the importance of neuronal glycogen in LTP. However, since only a subset of neurons was affected in this model, it remains unclear whether the differences between the GYS1^Nestin-KO^ and GYS1^Camk2a-KO^ lines are due to the lack of astrocytic glycogen and/or glycogen in another subtype of neuron.

In the present study, we generated a new mouse model, lacking GYS1 specifically in astrocytes (GYS1^Gfap-KO^). Our results clarify the specific contribution of astrocytic glycogen to cerebral function, confirming its role in synaptic plasticity and discarding its role in the prevention of epileptic seizures.

## 2 METHODS

### 2.1 Animals

Male and female mice aged 4 ± 1 months were used in this study. All experiments were carried out following European Union (2010/63/EU) and Spanish (BOE 34/11370-421, 2013) regulations for the use of laboratory animals. In addition, all experimental protocols were approved by the Ethics Committee of the Pablo de Olavide University. Animals were kept in collective cages (up to five animals per cage) on a 12-h light/dark cycle with constant temperature (21 ± 1°C) and humidity (50 ± 5%). After electrophysiological studies were initiated, mice were kept in individual cages until the end of the experiments. Animals were allowed access *ad libitum* to commercial mouse chow and water. The Gfap-Cre transgenic line 77.6 used in this study was purchased from Jackson laboratories (Stock #024098) and has been thoroughly characterized [12].

### 2.2 Biochemical analysis

Mice were euthanized by cervical dislocation and decapitation, and the brains were removed and hemisected. The cerebellum, hippocampus and cortex from each hemisphere were then dissected and frozen in liquid nitrogen. Samples were maintained at −80 ºC until use. Tissue lysates for Western blot were prepared as previously described [11] and sample protein content was determined by Bradford assay (BioRad). Lysates were loaded in 10% polyacrylamide gels and transferred to Immobilon membranes (Millipore) for Western blot. The following antibodies were used: anti-glycogen synthase (Cell Signaling cat# 3886) and anti-GFAP (Millipore cat# MAB360). The REVERT total protein stain was used as a loading control and densitometry was performed using Image Studio™ Lite (LI-COR BioSciences).

### 2.3 Animal preparation for the electrophysiological study

For electrode implantation, animals were anesthetized with 0.8–3% halothane from a calibrated Fluotec 5 (Fluotec-Ohmeda, Tewksbury, MA, USA) delivered via a homemade mask and vaporizer at a flow rate of 0.8 L/min oxygen. Briefly, animals were implanted with bipolar stimulating electrodes at the right Schaffer collaterals of the dorsal hippocampus and with a recording electrode in the ipsilateral CA1 area using stereotaxic coordinates [13]. Electrodes were made of 50 μm Teflon-coated tungsten wire (Advent Research Materials Ltd., Eynsham, England). The final location of the CA1 recording electrode was determined electrophysiologically, as described by some of us [11, 14]. Stimulating, recording and ground wires were soldered to a 6-pin socket. The socket was fixed to the skull with the help of three small screws and dental cement [11, 14].

### 2.4 Input/output curves and LTP procedures

Both input/output curves, paired-pulse facilitation (PPF), and LTP were evoked in behaving mice following procedures described elsewhere [11, 14]. For the electrophysiological study, the mouse was located in a small (5 × 5 × 5 cm) box, aimed to avoid over walking. For input/output curves, mice were stimulated at the CA3-CA1 synapse with single pulses of increasing intensities (0.02–0.4 mA). PPF was determined by applying double pulses with increasing inter-pulse intervals (10, 20, 40, 100, 200 and 500 ms) at a fixed intensity corresponding to ~40% of asymptotic values, as previously described [11]. Evoked field excitatory post-synaptic potentials (fEPSPs) were recorded with Grass P511 differential amplifiers, across a high impedance probe (2 × 10^12^ Ω; 10 pF), and with a bandwidth of 0.1 Hz-10 kHz (Grass-Telefactor, West Warwick, RI, USA).

For LTP measurements, baseline fEPSP values evoked at the CA3-CA1 synapse were collected 15 min prior to LTP induction using single 100 μs, square, biphasic pulses. Pulse intensity was set well below the threshold for evoking a population spike (0.15-0.25 mA); i.e., 30–40% of the intensity necessary for evoking a maximum fEPSP response [14, 15]. LTP was evoked with a high-frequency stimulus (HFS) protocol consisting of five 200 Hz, 100-ms trains of pulses at a rate of 1/s, repeated six times, at intervals of 1 min. The stimulus intensity during the HFS protocol was set at the same value as that used for generating baseline recordings to prevent the presentation of electroencephalographic seizures and/or large population spikes. After each HFS session, the same stimuli were presented individually every 20 s for 60 additional min and for 30 min on the following three days [11, 14]. Evoked fEPSPs were recorded as described above.

### 2.5 Induction of hippocampal seizures with kainate injections in implanted mice

Following procedures described elsewhere [16], we determined the propensity of control and GYS1^Gfap-KO^ mice to generate convulsive seizures in the hippocampal area. For this, we intraperitoneally (i.p.) administrated the AMPA/kainate receptor agonist kainate (8 mg/kg; Sigma, St. Louis, MO, USA) dissolved in 0.1 M phosphate buffered saline (PBS) pH = 7.4. Local field potentials and electrically evoked fEPSPs were recorded in the hippocampal CA1 area from 5 min before to 60 min after kainate injections.

### 2.6 Video monitoring of seizures after kainate injections

Animals were placed in individual cages and were administered with three consecutive i.p. injections of kainate (8 mg/kg per dose, 24 mg/kg total) one every 30 min from the onset of the experiment in order to induce convulsive non-lethal seizures. Seizure stages after kainate injections were evaluated as described previously [17–19]. After the first kainate injections, the animals developed hypoactivity and immobility (Stage I–II). After successive injections, hyperactivity (Stage III) and scratching (Stage IV) were often observed. Some animals progressed to a loss of balance control (Stage V) and further chronic whole-body convulsions (Stage VI). Extreme behavioural manifestations such as, uncontrolled hopping activity, or “popcorn behaviour” and continuous seizures (more than 1 minute without body movement control) were included in Stage VI. All behavioural assessments were performed blind to the experimental group (genotype) in situ, as well as recorded and reanalysed blind to the first analysis. Analysis consisted in the record of the time spent until the onset of the first seizure, the number of seizures per animal, the time spent on each grade, as well as the maximum grade reached by each animal.

### 2.7 Data collection and statistical analysis

fEPSPs and 1-V rectangular pulses corresponding to brain stimulation were stored digitally on a computer through an analog/digital converter (CED 1401 Plus, CED, Cambridge, England). Data were analyzed off-line for fEPSP recordings with the help of the Spike 2 (CED) program. Five successive fEPSPs were averaged, and the mean value of the amplitude (in mV) was determined. Computed results were processed for statistical analysis using the IBM SPSS Statistics 18.0 (IBM, Armonk, NY, United States). Data are represented as the mean ± SEM. Statistical significance of differences between groups was inferred by Two-way repeated measures ANOVA, followed by the Holm-Sidak method for all pairwise multiple comparison procedures. The Fisher exact test for data collected from kainate experiments. Statistical significance was set at *P* < 0.05. For the biochemical analyses and behavioral assessment of seizure susceptibility after kainate administration, computed results were processed for statistical analysis with PRISM 8.0 (GraphPAD Software, San Diego, USA). Data are represented as the mean ± SEM. Normality of the distributions was checked via the Shapiro–Wilk test; All tests performed were two-sided. Statistical significance of differences between groups was inferred by Student’s t-Test or Two-way ANOVA, followed by the Bonferroni post-hoc comparison for all pairwise multiple comparison procedures. Statistical significance was set at *P* < 0.05.

## 3 RESULTS

### 3.1 Generation of GYS1^Gfap-KO^ mice

GFAP is a cytoskeletal protein found in nearly all astrocytes and a common marker for this cell type. The astrocyte-specific inactivation of *Gys1* was achieved by crossing mice homozygous for the conditional *Gys1* allele [9] with mice expressing Cre recombinase under the control of the *Gfap* promoter [12]. Littermates that were homozygous for the conditional *Gys1* allele and negative for Cre recombinase expression were used as controls. To confirm the inactivation of *Gys1*, we measured GYS1 protein levels in cortex, hippocampus, and cerebellum by Western blot. GYS1 protein was greatly diminished in all regions (Fig. 1a). Quantification by densitometry showed that GYS1 expression was reduced by approximately 80% in the cortex and in the hippocampus, and 60% in the cerebellum (Fig. 1b). Total brain glycogen was decreased by more than 80% (Fig. 1c), which is consistent with the reduction in GYS1 protein. No changes in GFAP levels were observed (Fig. 1a, quantification not shown).

**Fig. 1.**
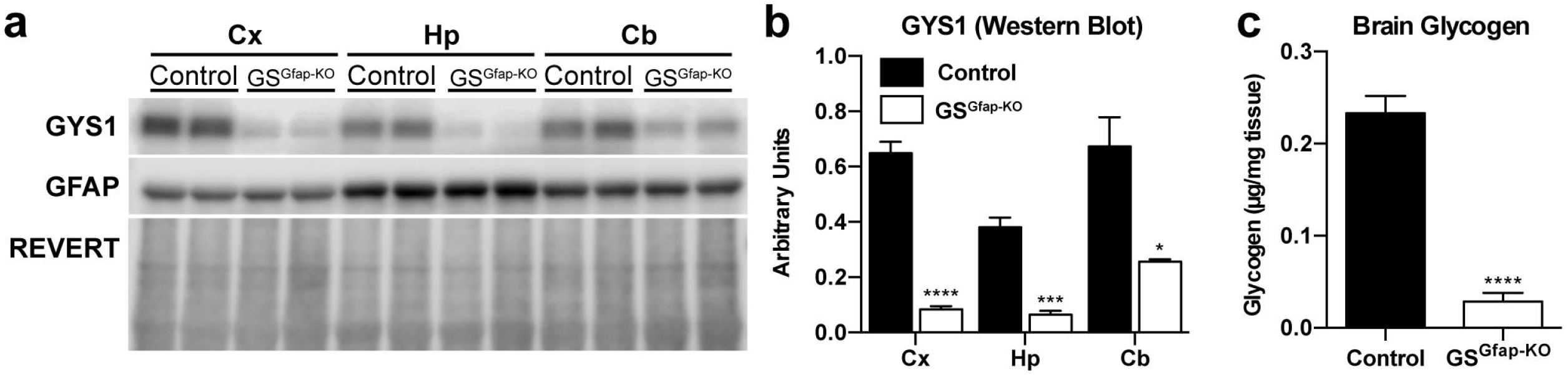
Analysis of GYS1, GFAP and glycogen levels in GS^Gfap-KO^ mice and controls. (a) Representative Western blot of GYS1 and GFAP protein levels in cortex (Cx), hippocampus (Hp) and cerebellum (Cb). REVERT protein stain (Li-COR BioSciences) was used as a loading control. (b) Quantification of GYS1 and GFAP protein levels by region normalized to total protein determined by REVERT. (c) Total brain glycogen in control versus GS^Gfap-KO^ animals. All data are expressed as average ±SEM (*n* = 4-6 per group). Significant differences were calculated using student’s t-test (∗, *P* < 0.05; ***, *P* < 0.001; ****, *P* < 0.0001).

### 3.2 Electrophysiological alterations at the CA3-CA1 synapse in GYS1^Gfap-KO^ mice

To study the consequences of the lack of astrocytic glycogen on synaptic function, we performed recordings of input/output curves, PPF and LTP evoked at the CA3-CA1 synapse of the hippocampus (Fig. 2a). In a first experimental step, we examined the response of CA1 pyramidal neurons to single pulses of increasing intensity (0.02–04 mA) presented to the ipsilateral Schaffer collaterals. Both control and GYS1^Gfap-KO^ mice presented similar increases in the amplitude of fEPSPs evoked at CA1 pyramidal neurons by the stimuli presented to Shaffer collaterals (Fig. 2b). These two input/output relationships were best fitted by sigmoid curves (*r* ≥ 0.9; *P* ≤ 0.001; not illustrated), suggesting the normal functioning of the CA3-CA1 synapse in both groups. However, the experimental group reached lower maximal fEPSP amplitudes than their littermate controls. No significant differences [Two-way repeated measures ANOVA; *F*_(19,266)_ = 1.222; *P* = 0.239] were observed overall between control and GYS1^Gfap-KO^ groups. However, fEPSPs evoked by three increasing intensities presented significant differences (All pairwise multiple comparison procedures; *P* < 0.05). We also performed an analysis of PPF at the CA3-CA1 synapse by applying double pulses at a fixed intensity with increasing inter-stimulus intervals (10, 20, 40, 100, 200, 500 ms). Both groups displayed facilitation at 20 and 40 ms intervals (Fig. 2c). GYS1^Gfap-KO^ exhibited no statistical difference in PPF compared to control animals [Two-way repeated measures ANOVA; *F*_(5,95)_ = 0.726; *P* = 0.606].

**Fig. 2.**
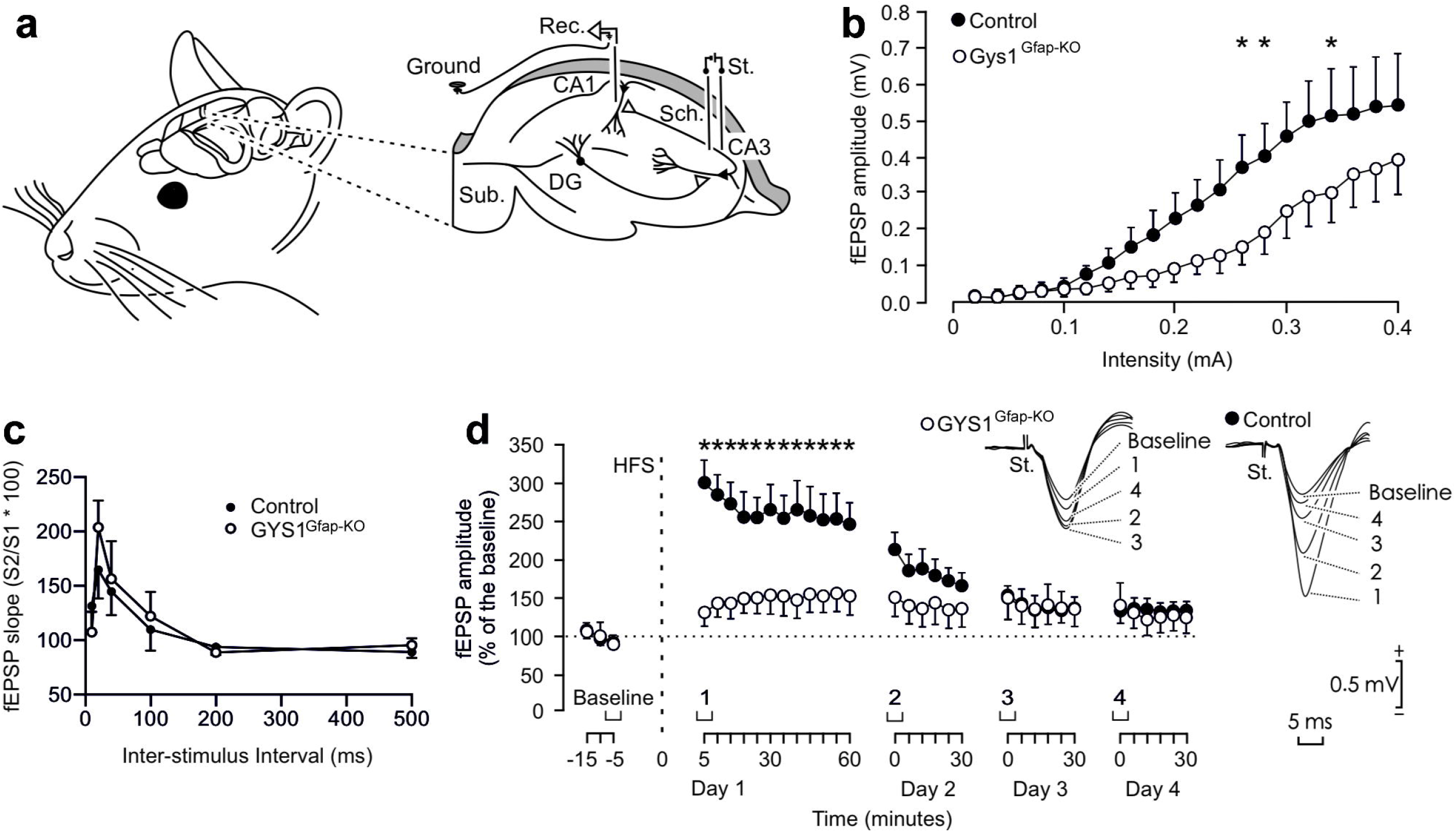
Electrophysiological properties of hippocampal synapses in behaving control and GYS1^Gfap-KO^ mice. (A) Animals were chronically implanted with bipolar stimulating (St.) electrodes in the right CA3 Schaffer collaterals and with a recording (Rec.) electrode in the ipsilateral CA1 area. DG, dentate gyrus; Sub., subiculum. (B) Input/output curves of fEPSPs evoked at the CA3-CA1 synapse via single pulses of increasing intensities (0.02–0.4 mA) in control and GYS1^Gfap-KO^ mice. Although no significant differences [Two-way repeated measures ANOVA; *F*_(19,266)_ = 1.222; *P* = 0.239] were observed between groups, fEPSPs evoked by three different intensities presented significant differences (All pairwise multiple comparison procedures; *P* < 0.05). (C) Paired-pulse facilitation in control and GYS1^Gfap-KO^ animals with increasing inter-stimulus intervals. No significant differences between the two groups were observed [Two-way repeated measures ANOVA; *F*_(5,95)_ = 1.222; *P* = 0.606]. (D) LTP evoked at the CA3-CA1 synapse of control and GYS1^Gfap-KO^ mice following the HFS session. The HFS was presented after 15 min of baseline recordings, at the time marked by the dashed line. LTP evolution was followed for four days. At the right are illustrated representative examples of fEPSPs collected from control and GYS1^Gfap-KO^ mice at the times indicated in the bottom graph. fEPSP amplitudes are given as a percentage of values measured from baseline recordings, and statistical differences between control and GYS1^Gfap-KO^ from two-way repeated measures ANOVA are shown (*, *P* ≤ 0.01). All data are expressed as average ± SEM (n= 7-9 mice/group).

In a following experimental step, we evoked LTP at the CA3-CA1 synapse of the two genotypes as an indication of long-term synaptic plasticity. It is well known that the hippocampus is involved in the acquisition of different types of associative [20, 21] and non-associative [22, 23] learning tasks and that the CA3-CA1 synapse is often selected for evoking LTP in behaving mice [14, 24]. For baseline values, animals were stimulated every 20 s for ≥ 15 min at the implanted Schaffer collaterals (Fig. 2d). Afterward, they were presented with a high frequency stimulus (HFS) protocol. Immediately after the HFS session, the same single stimulus used to generate baseline records was presented at the initial rate (3/min) for another 60 min. As illustrated in Fig. 2d, recording sessions were repeated for three additional days (30 min each). The control group presented a significant LTP when comparing baseline values with those collected following the HFS session (Holm-Sidak method, all pairwise multiple comparison procedures; *P* ≤ 0.041). Although the amplitude of fEPSPs also increased in GYS1^Gfap-KO^ mice following the HFS session, only a tendency was presented (*P* ≥ 0.671) (Fig. 2d). In addition, the amplitude of fEPSPs evoked in the control group was significantly [Two-way repeated measures ANOVA; *F*_(32,512)_ = 6.277; *P* < 0.001] larger and longer lasting than that evoked in the experimental group (Fig. 2d). In summary, GYS1^Gfap-KO^ mice display no change in PPF but a significantly impaired LTP compared to the littermate controls.

### 3.3 Seizure susceptibility in GYS1^Gfap-KO^ mice

Kainate is a widely used chemoconvulsant used to study seizure susceptibility in rodents [25]. We previously demonstrated that mice devoid of cerebral glycogen (GYS1^Nestin-KO^) are more susceptible to kainate-induced seizures [10]. To understand the participation of astrocytic glycogen in seizure susceptibility, we also assessed the response of GYS1^Gfap-KO^ mice to kainate-induced seizures in the pre-implanted animals. Evoked seizures in control and GYS1^Gfap–KO^ mice presented similar durations and profiles (Fig. 3a). Both groups presented a noticeable depression in the amplitude of evoked fEPSP recorded following a kainate-dependent seizure (Fig. 3b). Overall, there was no difference in the number of seizures observed per genotype (Fisher exact test; *P* = 0.657) (Fig. 3c).

**Fig. 3.**
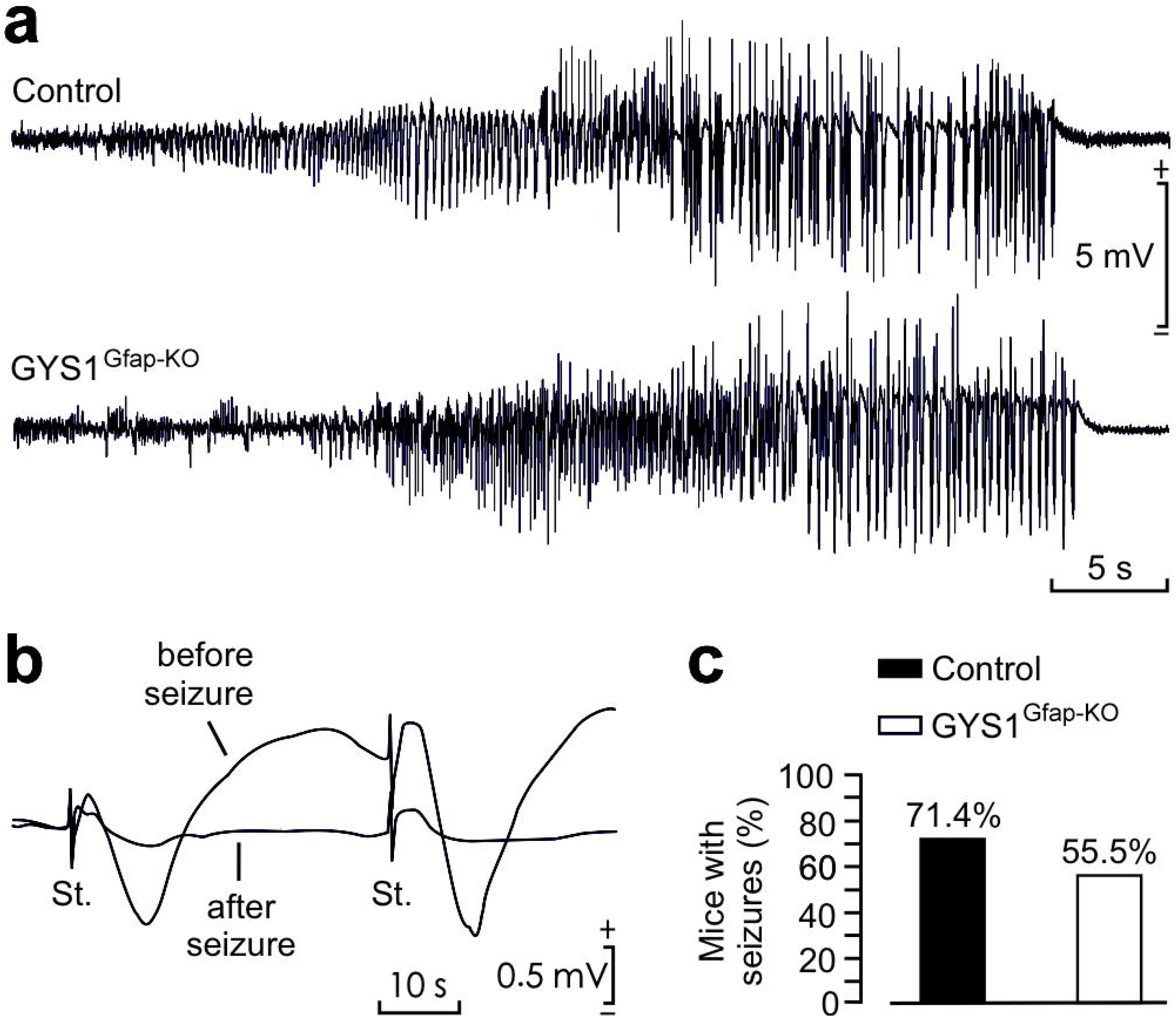
Kainate susceptibility of GYS1^Gfap-KO^ mice compared to controls. (a) Representative examples of hippocampal seizures evoked in control and GYS1^Gfap-KO^ mice following the administration of 8 mg/kg i.p. of kainate. (b) Representative examples of fEPSPs evoked before and immediately after a kainate-evoked seizure. (c) Percentage of control (*n* = 14) and GYS1^Gfap-KO^ (*n* = 9) mice presenting spontaneous seizures at the CA1 area during the recording period (60 min). No significant differences between groups (Fisher exact test; *P* = 0.657) were observed.

To corroborate these results, we employed an alternative seizure assessment protocol in a new cohort of mice. Mice were given three kainate injections (8 mg/kg, i.p. every 30 min) and video-recorded for 180 minutes to monitor their behavior (i.e. epileptic events) following the first injection. Mice from both genotypes reached similar severity stages (Fig. 4a). In the majority of the mice of both genotypes, seizures began approximately 15 minutes after the third dose of kainate (*P* = 0.9501, Student’s t-test; Fig. 4b). There were no significant differences between groups in the prioritary stage (the behavioral stage in which an animal spends the most time after kainate administration throughout the duration of the experiment) (*P* = 0.5441, Student’s t-test; Fig. 4c) nor the maximum stage (the most severe stage reached during the experiment) (*P* = 0.9644, Student’s t-test; Fig. 4d). Furthermore, there were no significant differences in the time spent per stage (Two-way ANOVA: Stage factor: *P* = 0.0007, Genotype factor: *P* = 0.2370; Fig. 4e) or in the number of seizures per animal after each injection (Two-way ANOVA: Administration factor: *P* < 0.0001, Genotype factor: *P* = 0.7980; Fig. 4f). These results unequivocally confirmed that GYS1^Gfap-KO^ animals present a similar seizure susceptibility compared to control littermates.

**Fig. 4.**
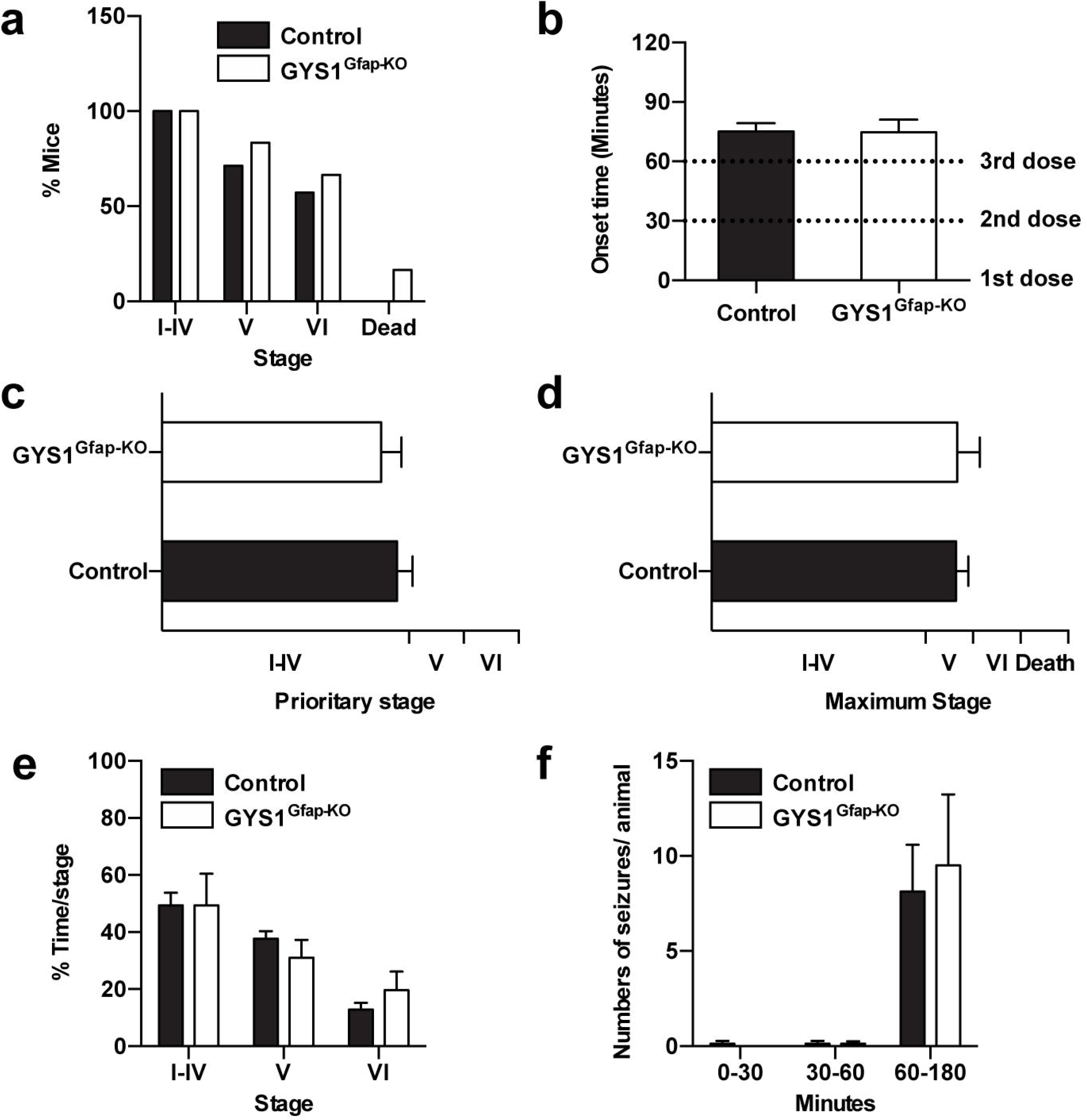
Comparison of kainate-induced seizure profile in control and GYS1^Gfap-KO^. 3-4 months old mice were subjected to three kainate injections (8 mg/kg every 30 min) and epileptic responses were analyzed for 180 minutes after the first injection. (a) Percentage of mice reaching seizure stages I to VI and kainate-induced mortality. (b) Onset of the epileptic activity. Student’s t-test (*P* = 0.9501). (c) Prioritary stage displayed by each animal during the course of the experiment. Student’s t-test (*P* = 0.5441). (d) Maximum stage reached by each animal during the course of the experiment. Student’s t-test (*P* = 0.9644). (e) Percentage of time spent on each stage during the course of the experiment. Two-way ANOVA (Stage factor: *P* = 0.0007; Genotype factor: *P* = 0.2370). (f) Number of seizures experimented per animal divided on time segments after the first, second and third kainate administrations. Two-way ANOVA (Administration factor: *P* < 0.0001; Genotype factor: *P* = 0.7980). All data are expressed as average ± SEM (*n* = 6-7 mice/group).

## 4 DISCUSSION

In this study, we analyze for the first time the physiological consequences of removing glycogen specifically from astrocytes by means of transgenic tools. By comparing this mouse to previous models lacking glycogen in the entire CNS [9, 10] or only in Camk2a-positive excitatory neurons of the forebrain [11], we are able to define the specific physiological roles of glycogen in astrocytes versus neurons.

Using Cre/Lox technology, we deleted *Gys1* in GFAP-positive cells to eliminate only astrocytic glycogen synthesis. Western blot analyses confirmed a clear reduction in GYS1 protein levels in hippocampus, cortex and cerebellum of GYS1^Gfap-KO^ mice (Fig. 1, a and b). Total brain glycogen content was also greatly reduced relative to littermate controls (Fig. 1c). These results are consistent with the well-documented observation that the majority of brain glycogen is stored in astrocytes, but also point to a remaining significant fraction of non-astrocytic GYS1 expression and glycogen synthesis, likely in neurons.

We also studied synaptic function in the GYS1^Gfap-KO^ model via stimulation of the CA3-CA1 synapse in the hippocampus. Input/output curves showed no overall significant difference between groups, but at some intensities, evoked fEPSPs were statistically higher in the control group compared to the GYS1^Gfap-KO^ animals, and the latter group reached a lower maximum fEPSP (Fig. 2b). These results suggest that the absence of astrocytic glycogen may reduce basal synaptic strength. A similar situation is obtained with inhibitors of astrocytic glutamate transport, which also reduce basal EPSPs since the accumulation of glutamate causes presynaptic inhibition [26]. Astrocytes take up synaptic glutamate and convert it to glutamine for transfer to neurons, where it is recycled into glutamate and repackaged into synaptic vesicles, a process known as the glutamate/glutamine cycle [27, 28]. Since astrocytic glycogen has been shown to play a role in both glutamate uptake and recycling [29, 30], the lack of astrocytic glycogen could cause impaired glutamate uptake leading to presynaptic inhibition and therefore lower synaptic strength. While PPF experiments with increasing inter-pulse intervals showed significantly greater facilitation in GYS1^Nestin-KO^ mice [10], in the present study, GYS1^Gfap-KO^ mice only lack astrocytic glycogen displayed only a trend toward increased PPF, with no statistical difference (Fig. 2c). PPF is typically attributed to presynaptic mechanisms such increased [Ca^2+^] in the presynaptic terminal [31]. The altered PPF observed in GYS1^Nestin-KO^ mice could be exclusively neuronal in origin.

We also observed an impairment in hippocampal LTP in GYS1^Gfap-KO^ animals (Fig. 2d). However, the LTP impairment in the GYS1^Gfap-KO^ animals was not as pronounced as was observed in the GYS1^Nestin-KO^ mice [9]. The present results are reminiscent of those from the GYS1^Camk2a-KO^ mice, in which LTP was still observed, although it was significantly impaired [11]. Collectively, these three mouse models demonstrate that both astrocytic and neuronal glycogen contribute to LTP. Previous studies using pharmacological agents have shown that astrocytic glycogen is important for long-term, but not short-term, memory formation [32, 33]. The presence of a normal PPF, a measure of short-term plasticity, and the major impairment in LTP that we observed in the GYS1^Gfap-KO^ model are consistent with these observations.

The most surprising result regarding the GYS1^Gfap-KO^ line is its susceptibility to kainate-induced epilepsy. We previously showed that GYS1^Nestin-KO^ mice were more susceptible to seizures induced by a single convulsive dose of kainate (8mg/kg, i.p.) [10]. However, using the same protocol, we detected no statistical difference in the GYS1^Gfap-KO^ line (Fig. 3). Utilizing a second experimental protocol with three consecutive doses of kainate (8mg/kg, i.p., every 30 minutes) we found no statistical differences between the groups in the seizure stages achieved (Fig. 4a), seizure onset (Fig. 4b), prioritary or maximum stage reached (Fig. 4, c and d), time spent per stage (Fig. 4e) or number of seizures per animal (Fig. 4f). These results unequivocally show that GYS1^Gfap-KO^ animals are not more susceptible to kainate than their littermate controls. Altered glycogen metabolism has been linked to seizures, as reviewed elsewhere [34–37]. A commonly held view is that altered astrocytic glycogen metabolism induces neuronal excitability via impaired glutamate and K^+^ uptake. However, herein we show mice lacking astrocytic glycogen do not have more kainate-induced seizures. Since GYS1^Camk2-KO^ mice also show unaltered kainate susceptibility [11], collectively these mouse models suggest that seizure susceptibility in the GYS1^Nestin-KO^ line is a consequence of the lack of glycogen in another cell type. Inhibitory neurons play a critical role in suppressing excitability, and their dysfunction is associated with epilepsy in rodent models and humans [38]. Therefore, our results suggest that glycogen in inhibitory neurons might be critical for their regulatory role. This possibility will be addressed in future studies.

In summary, the GYS1^Gfap-KO^ mouse model illustrates the specific contribution of astrocytic glycogen to the physiological roles of glycogen in the brain, further clarifying how brain glycogen is involved in memory and epilepsy. Our results confirm that astrocytic glycogen plays an active role in long-term synaptic plasticity. However, the lack of astrocytic glycogen does not increase susceptibility to kainate-induced seizures in these mice. These data point to a role of neuronal glycogen in cerebral functions, most importantly in the regulation of excitability. A thorough understanding of these processes is essential for better management and treatment of neurological disorders.

## AUTHOR CONTRIBUTIONS

JD and JJG conceived the study. JD generated and maintained the GYS1^Gfap-Cre^ line. JD and MKB collected brain tissues and performed biochemical analyses. AG and JMD-G performed electrophysiological studies before and after single kainate injections. AH and JAR performed seizure video-monitoring with multiple kainate injections. All authors analyzed data and contributed to the writing of the manuscript.

## ACKNOWLEDGEMENTS

We thank Anna Adrover, Emma Veza, Anna Guitart and the IRB Histopathology Facility for technical assistance and Laura Alcaide, Vanessa Hernandez, María Sánchez Enciso and José M. González Martín for their help in animal handling and care. We also thank Olga Varea for helpful advice and discussions.

IRB Barcelona and IBEC are recipients of a Severo Ochoa Award of Excellence from MINECO (Government of Spain). This study was supported by grants from the MINECO (BFU2017-82375-R to AG and JMD-G, RTI2018-099773-B-I00 to JADR and AH and BFU2017-84345-P to JD and JG), the CIBER de Diabetes y Enfermedades Metabólicas Asociadas (ISCIII, Ministerio de Ciencia e Innovación), and a grant from the National Institutes of Health (NIH NINDS P01NS097197) to JG. The project also received funding from “la Caixa” Foundation (ID 100010434) under the agreement LCF/PR/HR19/52160007 with JADR. MKB has received funding from the European Union’s Horizon 2020 research and innovation program under the Marie Skłodowska-Curie grant agreement No. 754510.

## REFERENCES

1. Brown AM (2004) Brain glycogen re-awakened. J Neurochem 89:537–552. https://doi.org/10.1111/j.1471-4159.2004.02421.x

2. Saez I, Duran J, Sinadinos C, et al (2014) Neurons Have an Active Glycogen Metabolism that Contributes to Tolerance to Hypoxia. J Cereb Blood Flow Metab 34:945–955. https://doi.org/10.1038/jcbfm.2014.33

3. Rubio-Villena C, Viana R, Bonet J, et al (2018) Astrocytes: new players in progressive myoclonus epilepsy of Lafora type. Human Molecular Genetics 27:1290–1300. https://doi.org/10.1093/hmg/ddy044

4. Gibbs ME (2016) Role of Glycogenolysis in Memory and Learning: Regulation by Noradrenaline, Serotonin and ATP. Front Integr Neurosci 9:. https://doi.org/10.3389/fnint.2015.00070

5. Alberini CM, Cruz E, Descalzi G, et al (2018) Astrocyte glycogen and lactate: New insights into learning and memory mechanisms. Glia 66:1244–1262. https://doi.org/10.1002/glia.23250

6. Duran J, Guinovart JJ (2015) Brain glycogen in health and disease. Molecular Aspects of Medicine 46:70–77. https://doi.org/10.1016/j.mam.2015.08.007

7. Oe Y, Akther S, Hirase H (2019) Regional Distribution of Glycogen in the Mouse Brain Visualized by Immunohistochemistry. Adv Neurobiol 23:147–168. https://doi.org/10.1007/978-3-030-27480-1_5

8. Schulz A, Sekine Y, Oyeyemi MJ, et al (2020) The stress-responsive gene GDPGP1/mcp-1 regulates neuronal glycogen metabolism and survival. J Cell Biol 219:. https://doi.org/10.1083/jcb.201807127

9. Duran J, Saez I, Gruart A, et al (2013) Impairment in Long-Term Memory Formation and Learning-Dependent Synaptic Plasticity in Mice Lacking Glycogen Synthase in the Brain. J Cereb Blood Flow Metab 33:550–556. https://doi.org/10.1038/jcbfm.2012.200

10. López-Ramos JC, Duran J, Gruart A, et al (2015) Role of brain glycogen in the response to hypoxia and in susceptibility to epilepsy. Front Cell Neurosci 9:. https://doi.org/10.3389/fncel.2015.00431

11. Duran J, Gruart A, Varea O, et al (2019) Lack of Neuronal Glycogen Impairs Memory Formation and Learning-Dependent Synaptic Plasticity in Mice. Front Cell Neurosci 13:374. https://doi.org/10.3389/fncel.2019.00374

12. Gregorian C, Nakashima J, Le Belle J, et al (2009) Pten Deletion in Adult Neural Stem/Progenitor Cells Enhances Constitutive Neurogenesis. Journal of Neuroscience 29:1874–1886. https://doi.org/10.1523/JNEUROSCI.3095-08.2009

13. Franklin KBJ, Paxinos G (2008) The mouse brain in stereotaxic coordinates, 3. ed. Elsevier, AP, Amsterdam

14. Gruart A (2006) Involvement of the CA3-CA1 Synapse in the Acquisition of Associative Learning in Behaving Mice. Journal of Neuroscience 26:1077–1087. https://doi.org/10.1523/JNEUROSCI.2834-05.2006

15. Gureviciene I, Ikonen S, Gurevicius K, et al (2004) Normal induction but accelerated decay of LTP in APP + PS1 transgenic mice. Neurobiology of Disease 15:188–195. https://doi.org/10.1016/j.nbd.2003.11.011

16. Valles◻Ortega J, Duran J, Garcia◻Rocha M, et al (2011) Neurodegeneration and functional impairments associated with glycogen synthase accumulation in a mouse model of Lafora disease. EMBO Mol Med 3:667–681. https://doi.org/10.1002/emmm.201100174

17. Carulla P, Bribián A, Rangel A, et al (2011) Neuroprotective role of PrP ^C^ against kainate-induced epileptic seizures and cell death depends on the modulation of JNK3 activation by GluR6/7–PSD-95 binding. MBoC 22:3041–3054. https://doi.org/10.1091/mbc.e11-04-0321

18. Rangel A, Madroñal N, Massó AG i., et al (2009) Regulation of GABAA and Glutamate Receptor Expression, Synaptic Facilitation and Long-Term Potentiation in the Hippocampus of Prion Mutant Mice. PLoS ONE 4:e7592. https://doi.org/10.1371/journal.pone.0007592

19. Rangel A, Burgaya F, Gavín R, et al (2007) Enhanced susceptibility of Prnp-deficient mice to kainate-induced seizures, neuronal apoptosis, and death: Role of AMPA/kainate receptors. Journal of Neuroscience Research 85:2741–2755. https://doi.org/10.1002/jnr.21215

20. Thompson RF (2005) In Search of Memory Traces. Annu Rev Psychol 56:1–23. https://doi.org/10.1146/annurev.psych.56.091103.070239

21. Gruart A, Leal-Campanario R, López-Ramos JC, Delgado-García JM (2015) Functional basis of associative learning and its relationships with long-term potentiation evoked in the involved neural circuits: Lessons from studies in behaving mammals. Neurobiology of Learning and Memory 124:3–18. https://doi.org/10.1016/j.nlm.2015.04.006

22. Clarke JR, Cammarota M, Gruart A, et al (2010) Plastic modifications induced by object recognition memory processing. Proceedings of the National Academy of Sciences 107:2652–2657. https://doi.org/10.1073/pnas.0915059107

23. Moser EI, Moser M-B, McNaughton BL (2017) Spatial representation in the hippocampal formation: a history. Nat Neurosci 20:1448–1464. https://doi.org/10.1038/nn.4653

24. Bliss TV, Collingridge GL (2013) Expression of NMDA receptor-dependent LTP in the hippocampus: bridging the divide. Mol Brain 6:5. https://doi.org/10.1186/1756-6606-6-5

25. Lévesque M, Avoli M (2013) The kainic acid model of temporal lobe epilepsy. Neuroscience & Biobehavioral Reviews 37:2887–2899. https://doi.org/10.1016/j.neubiorev.2013.10.011

26. Oliet SHR (2001) Control of Glutamate Clearance and Synaptic Efficacy by Glial Coverage of Neurons. Science 292:923–926. https://doi.org/10.1126/science.1059162

27. McKenna MC (2007) The glutamate-glutamine cycle is not stoichiometric: Fates of glutamate in brain. Journal of Neuroscience Research 85:3347–3358. https://doi.org/10.1002/jnr.21444

28. Bak LK, Schousboe A, Waagepetersen HS (2006) The glutamate/GABA-glutamine cycle: aspects of transport, neurotransmitter homeostasis and ammonia transfer. Journal of Neurochemistry 98:641–653. https://doi.org/10.1111/j.1471-4159.2006.03913.x

29. Gibbs ME, Lloyd HGE, Santa T, Hertz L (2007) Glycogen is a preferred glutamate precursor during learning in 1-day-old chick: Biochemical and behavioral evidence. Journal of Neuroscience Research 85:3326–3333. https://doi.org/10.1002/jnr.21307

30. Schousboe A, Sickmann HM, Walls AB, et al (2010) Functional Importance of the Astrocytic Glycogen-Shunt and Glycolysis for Maintenance of an Intact Intra/Extracellular Glutamate Gradient. Neurotox Res 18:94–99. https://doi.org/10.1007/s12640-010-9171-5

31. Zucker RS, Regehr WG (2002) Short-Term Synaptic Plasticity. Annu Rev Physiol 64:355–405. https://doi.org/10.1146/annurev.physiol.64.092501.114547

32. Suzuki A, Stern SA, Bozdagi O, et al (2011) Astrocyte-Neuron Lactate Transport Is Required for Long-Term Memory Formation. Cell 144:810–823. https://doi.org/10.1016/j.cell.2011.02.018

33. Gibbs ME, Anderson DG, Hertz L (2006) Inhibition of glycogenolysis in astrocytes interrupts memory consolidation in young chickens. Glia 54:214–222. https://doi.org/10.1002/glia.20377

34. Bak LK, Walls AB, Schousboe A, Waagepetersen HS (2018) Astrocytic glycogen metabolism in the healthy and diseased brain. J Biol Chem 293:7108–7116. https://doi.org/10.1074/jbc.R117.803239

35. DiNuzzo M, Mangia S, Maraviglia B, Giove F (2015) Does abnormal glycogen structure contribute to increased susceptibility to seizures in epilepsy? Metab Brain Dis 30:307–316. https://doi.org/10.1007/s11011-014-9524-5

36. DiNuzzo M, Mangia S, Maraviglia B, Giove F (2014) Physiological bases of the K+ and the glutamate/GABA hypotheses of epilepsy. Epilepsy Research 108:995–1012. https://doi.org/10.1016/j.eplepsyres.2014.04.001

37. Duran J, Gruart A, López-Ramos JC, et al (2019) Glycogen in Astrocytes and Neurons: Physiological and Pathological Aspects. In: DiNuzzo M, Schousboe A (eds) Brain Glycogen Metabolism. Springer International Publishing, Cham, pp 311–329

38. Maglóczky Z, Freund TF (2005) Impaired and repaired inhibitory circuits in the epileptic human hippocampus. Trends in Neurosciences 28:334–340. https://doi.org/10.1016/j.tins.2005.04.002

